# What drives cultural ecosystem services in mountain protected areas? An AI-assisted answer using social media

**DOI:** 10.1101/2025.07.23.666097

**Authors:** José Carlos Pérez-Girón, Carlos Javier Navarro, Akram Elghouat, Rohaifa Khaldi, Domingo López, Salvador Arenas-Castro, Ana del Águila, Ricardo Moreno-Llorca, Nuria Pistón, Luis F. Romero, Ana Sofía Vaz, Siham Tabik, Javier Martínez-López, Domingo Alcaraz-Segura

## Abstract

High mountain protected areas (PAs) are increasingly recognized not only for their role in conserving biodiversity but also for their contribution to the provision of cultural ecosystem services (CES). Despite their relevance, CES remain underrepresented in conservation planning, particularly due to challenges in quantifying their spatial distribution. This study combines geolocated social media data and ecological niche models (ENMs) to assess the spatial patterns and key drivers of CES supply across eight mountain PAs spanning distinct biogeographical regions in Spain and Portugal. Using deep learning techniques to classify more than 200,000 photographs into ten CES categories, we evaluated model performance under two modeling approaches and identified the most influential environmental and social predictors. Most CES categories exhibited good model performance (Boyce index > 0.5), though variation existed across services and regions. Nature & Landscape and Gastronomy CES showed strong associations with park boundaries and human settlements, respectively, while Religious and Cultural CES were spatially linked to culturally significant landmarks.. Our findings demonstrate the potential of combining social media data with ENMs to map CES distributions and reveal both universal and context-dependent drivers. This approach offers valuable insights for integrating CES into PA management and spatial planning, supporting more holistic and culturally inclusive conservation strategies.

## 1. Introduction

In the Anthropocene, human-driven environmental transformations are accelerating at an unprecedented rate, disrupting ecosystems and jeopardizing the long-term conservation of biodiversity and the essential ecosystem services that sustain human well-being (IPBES, 2019; MEA, 2005). High mountain ecosystems, often considered isolated and resilient, are on the frontline of this battle, due to their high level of sensitivity to climate change (Kohler et al., 2010). Over a century ago, the declaration of the world’s first National Parks symbolized the beginning of the collective efforts to protect and conserve nature. Since then, the different geopolitical entities have developed conservation policies, management strategies, and legislative frameworks to safeguard Protected Areas (PAs). These include different protection figures and levels such as UNESCO Biosphere Reserves, the Natura 2000 network, and National Parks between others, all of which play a key role in protecting biodiversity, preserving ecosystems, and subsequently, supporting the provision of ecosystem services. A provisioning ability that occurs not only within its boundaries but also in its surrounding areas (Castro et al., 2015; Palomo et al., 2013). However, while the primary focus of conservation efforts has historically been on biodiversity conservation and the preservation of natural landscapes, the recognition of ecosystem services values into PAs has been recently added (Pu et al., 2023). An example of this is the European Union’s prioritization of nature conservation and the capacity to provide ecosystem services as part of its commitment to the Sustainable Development Goals (Xu and Peng, 2024). Despite these advances, significant knowledge gaps remain, particularly regarding how people perceive, value, and prioritize specific locations for cultural experiences in mountain PAs. This is especially relevant for Cultural Ecosystem Services (CES), which have received comparatively less attention. As highlighted by Schirpke et al., (2021), most research to date has emphasized the importance of mountain PAs in delivering provisioning and regulating services, while CES remains understudied.

Cultural Ecosystem Services (CES) are defined as the non-material benefits that people derive from ecosystems, which influence their physical and mental well-being, including recreation, spiritual fulfillment and aesthetic appreciation, among others (Haines-Young and Potschin-Young, 2018; MEA, 2005). Beyond their intrinsic value, CES play a fundamental role in sustaining tourism revenues, shaping cultural heritage and traditions, and fostering environmental behaviors that support nature conservation. On the other hand, CES can also have unintended consequences for biodiversity conservation, particularly when the overuse of natural areas leads to habitat degradation, wildlife disturbance, or conflicts between conservation goals and human activities (Belsoy et al., 2012; Penteriani et al., 2017; Spalding et al., 2023; Taff et al., 2019). These trade-offs underscore the need to understand the factors that motivate people to choose specific places to experience CES, as this knowledge is essential for balancing cultural benefits with ecological sustainability. Incorporating CES into the design and management of PAs can help balance conservation objectives with human well-being, ensuring that both nature and people benefit from sustainable interactions with these landscapes (Portman, 2013).

Quantifying CES presents significant challenges given their subjective nature and context-dependent, intangibility, spatio-temporal variability or lack of data, making them difficult to standardize and evaluate (Willcock et al., 2021). For that, their assessment typically relies on qualitative data, social surveys, participatory mapping, geospatial analysis of visitation patterns (Felipe-Lucia et al., 2024; Rodríguez-Morales et al., 2020). Recently, the emergence of social media platforms has revolutionized the way CES are mapped and analyzed (Cardoso et al., 2022; Moreno-Llorca et al., 2020; Muñoz et al., 2020; Navarro et al., 2025; Vaz et al., 2020, 2019). Geolocated data from social media, such as photographs, provide valuable insights into people’s interactions with natural landscapes, offering a cost-effective and large-scale approach to quantifying CES supply and demand (Oteros-Rozas et al., 2018; Tenkanen et al., 2017). However, the use of social media photographs to evaluate CES presents several challenges (Ghermandi et al., 2023; Oteros-Rozas et al., 2018; Schirpke et al., 2023; Sinclair et al., 2023). First, they are spatially biased toward accessible locations, as users tend to capture and share photos from well-known, aesthetically appealing, and easily reachable sites, leading to an overrepresentation of certain sites while more remote or less visually striking areas remain underrepresented. Second, they are content biased, since users select photograph subjects that align with personal interests, trends, or platform-specific engagement patterns, while repeated uploading of multiple photos of the same subject further amplify these biases. Lastly, social media photographs indicate the presence of certain CES but they lack information about areas where these services are absent. All together, combined with the inherent uncertainty of geolocated coordinates, can hinder accurate spatial analysis and lead to an incomplete understanding of CES distribution.

Despite the increasing use of social media data to assess CES, our understanding of the key environmental and social drivers that shape people’s preferences across different types of services, types of protected areas, or biogeographical regions remains limited. Most studies have focused on identifying areas of CES supply or demand, mapping hotspots, or analyzing patterns of spatial visitation (Da Silva et al., 2024; Lingua et al., 2023; Schirpke et al., 2018a). While these approaches have been valuable in showing where people interact with nature, they often overlook the deeper question of why certain places are consistently valued across different landscapes, cultures, or CES types (Havinga et al., 2022; Vaz et al., 2020). For instance, symbolic natural and cultural features, such as peaks, lakes, charismatic wildlife, or historic landmarks, are commonly assumed to influence how people perceive, experience and value nature (Rüdisser et al., 2019; Schirpke et al., 2018b). Similarly, social and cultural factors, including traditions, recreational trends, or even digital behaviors, influence how individuals express their preferences on social media platforms. However, the extent to which these drivers act universally across CES categories, spatial contexts, or protected areas (e.g., National Parks vs Natura 2000 sites) remains largely unknown. To address this complexity, a broad range of environmental factors characterizing the structure, composition, and dynamics of social-ecological systems are commonly selected for modeling CES, incorporating both biophysical and socio-cultural dimensions (Navarro et al., 2025; Vaz et al., 2020).

To bridge the gap between observed human-nature interactions and their spatial drivers, an ecological modeling perspective can offer valuable insights. In this context, classified social media photographs provide geolocated occurrences of CES, which serve as the foundation (or presences) for applying ecological niche models (ENMs). Originally developed for biodiversity research, ENMs are increasingly being adapted to study cultural interactions with landscapes (Clemente et al., 2019; Navarro et al., 2025). By modeling the environmental and spatial conditions associated with CES presence, ENMs create mathematical representations of the CES niche, helping to identify suitable areas for each CES and project them onto geographic space (Sillero et al., 2021). Therefore, this approach provides a powerful framework for uncovering the spatial drivers of CES and identifying potential areas of high cultural value.

Our goal is to enhance our understanding of how environmental and social factors shape CES distributions in mountain protected areas across biogeographical regions. For this, we addressed the next questions:

1. Can social media data and deep learning models be used to provide a reliable source of presence data to accurately map the supply of CES with ecological niche models?
2. Are there any universal environmental and social drivers that consistently promote the supply of CES independently of CES type, the type of protected area, or the biogeographical region? More specifically:

2.1. Are certain drivers consistently associated with CES supply across different CES types, or is each CES type driven by different environmental and social factors?
2.2. Are these drivers consistent across different spatial contexts, or do they vary across parks or biogeographical regions?
3. Does the category of protected area influence both the suitability of each CES type and the overall richness of CES?

## 2. Material and methods

### 2.1. Study area and protection levels

Given the high sensitivity of high mountain areas to the effects of climate change, the study area comprises the eight mountain National Parks located in Portugal and Spain, spanning three distinct biogeographical regions: Teide (in the Macaronesia region), Peneda-Gerês, and Picos de Europa (in the Atlantic region), Ordesa y Monte Perdido and Aigüestortes y Estany de Sant Maurici (in the Alpine region), and Sierra Nevada, Guadarrama and Sierra de las Nieves (in the Mediterranean region) (Figure 1a). Recognizing that protected areas are influenced by their surrounding landscapes, the study area encompasses the extension of their Socioeconomic Influence Areas, which are defined as the territories comprising the municipalities contributing land to the park and, in exceptional cases, additional areas directly linked to the park as specified in the corresponding declaratory legislation. This area was expanded to include all surrounding municipalities in a 100 m buffer.

**Figure 1.**
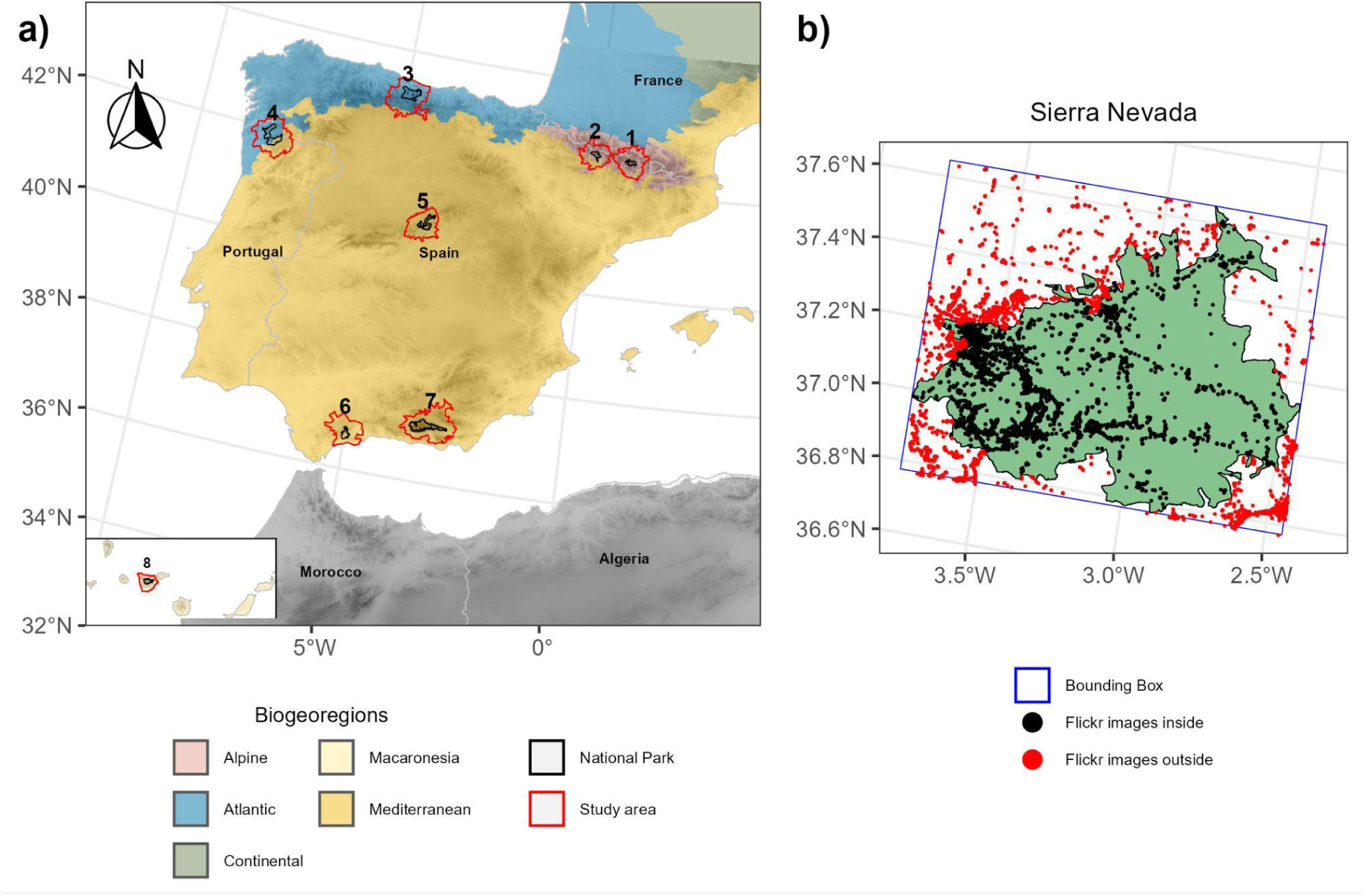
a) European biogeographical regions and location of the eight mountain National Parks across the Iberian Peninsula and the Canary Islands. The National Parks included are: (1) Aigüestortes i Estany de Sant Maurici, (2) Ordesa y Monte Perdido, (3) Picos de Europa, (4) Peneda-Gerês, (5) Sierra de Guadarrama, (6) Sierra de las Nieves, (7) Sierra Nevada, and (8) Teide. b) Example of Flickr image downloads within the bounding box of Sierra Nevada, followed by data cleaning procedures based on geolocation.

However, within these study areas, national parks coexist with other protection designations, including biosphere reserves and areas classified under the Natura 2000 network. To integrate these different protection levels, their spatial boundaries were obtained, reprojected into the EPSG:3035 coordinate system, and rasterized. Subsequently, all protection layers were intersected to generate a binary classification system representing the protection level of each grid cell. In this system, the first position indicates whether the cell is within a national park, the second denotes inclusion within a Natura 2000 site, and the third specifies whether the cell falls within a biosphere reserve. For example, a code of ‘110’ indicates that the cell is protected under both a national park and a Natura 2000 site but is not part of a biosphere reserve. A summary of the area occupied by each protection level within each study area is presented in Table S1.

### 2.2. CES occurrence data

All publicly shared geotagged photographs from 2015 to 2022 within the bounding boxes of each National park and their corresponding socioeconomic influence areas, a total of 338,259 images, were retrieved from the Application Programming Interface (API) of Flickr in July 2023 and subsequently downloaded. Metadata recorded for each image included the unique user identifier, photograph ID, geographic coordinates (latitude and longitude), date and time of capture and a URL linking the photo to the Flickr website, among others. After downloading, corrupted images and those from Flickr that fell outside of the study area were removed (Figure 1b). Following this filtering process, a total of 241,582 Flickr images remained.

Filtered images were labeled following a semi-automatic process based on two unsupervised and supervised approaches. Initially, features were extracted from 241,582 filtered images using the pre-trained large vision transformer model DINOv2 (Oquab et al., 2024) to generate high-quality image embeddings. Subsequently, dimensionality reduction was performed on these embeddings utilizing the Uniform Manifold Approximation and Projection (UMAP) algorithm (McInnes et al., 2020). The resulting low-dimensional vectors were clustered into 300 groups based on visual similarity using the k-means algorithm. A representative subset of images nearest to each cluster centroid was manually labeled, after which labels were propagated to the remaining images within each cluster, significantly reducing manual annotation efforts while preserving label consistency. Following this propagation step, a stratified and representative subset consisting of 7,110 images was further visualized and carefully reviewed by three independent domain experts to verify and correct labels as needed. This refined set of images served as the Ground Truth dataset for subsequent supervised methods.

In the first supervised approach, we performed fine-tuning of the pre-trained DINOv2 models (both small and base variants) utilizing this Ground Truth subset with an 80/20 train/test partition. The supervised approach involved two sequential steps, involving Linear Probing (batch size = 128, epochs = 5, learning rate = 1e-3) to adapt the classification head to our task, followed by full model Fine-Tuning (batch size = 128, epochs = 5, learning rate = 1e-6, following recommendations from Oquab et al., 2024) in order to reduce underperforming in future Out Of Distribution subsets (Kumar et al., 2022). Early stopping was applied in both stages to prevent overfitting and enhance generalization performance. Additionally, various architectural strategies were explored, with the final optimal solution involving the addition of a classification head comprising two fully connected neural network layers activated by ReLU functions. Finally, the full fine-tuning procedure, using the complete Ground Truth dataset, yielded a robust classifier, which was subsequently utilized for inference on the remaining unlabeled images up to the full dataset size of 241,582 images.

Similarly, images were classified using a zero-shot classification approach based on prompt engineering with OPENAI’s ChatGPT (GPT-4.1 model). Labeling was performed using the predefined categories described below. Eight candidate prompts were evaluated by comparing their classification results with those of the expert-labelled Ground Truth subset. The prompt with the highest macro-averaged F1 score was selected to ensure the best agreement with the annotations of the field experts. This optimised prompt was then applied to the entire image dataset.

Both approaches classified the filtered images into 10 CES-related classes, which were defined based on previous research and expertise (see Table S2) (Moreno-Llorca et al., 2020), and a discard category called Not relevant. Only images for which both approaches produced consistent labels were retained for further analysis.

A random subset of 1082 images covering all parks, municipalities, and labels, was manually and independently annotated to evaluate the performance of CES classification and niche models (Table 1). Each image was assigned up to three CES categories from the ten previously defined classes or labeled as non-relevant. This manually labeled subset was excluded from the training dataset to ensure independent validation. The quality of CES labels in the training dataset was assessed using the F1-score for each category (Table S3). Categories with an F1-score below 0.6 were excluded from the modeling process due to their high misclassification rates. As a result, the categories Sports and Sun & Beach were removed from further analyses.

**Table 1.**
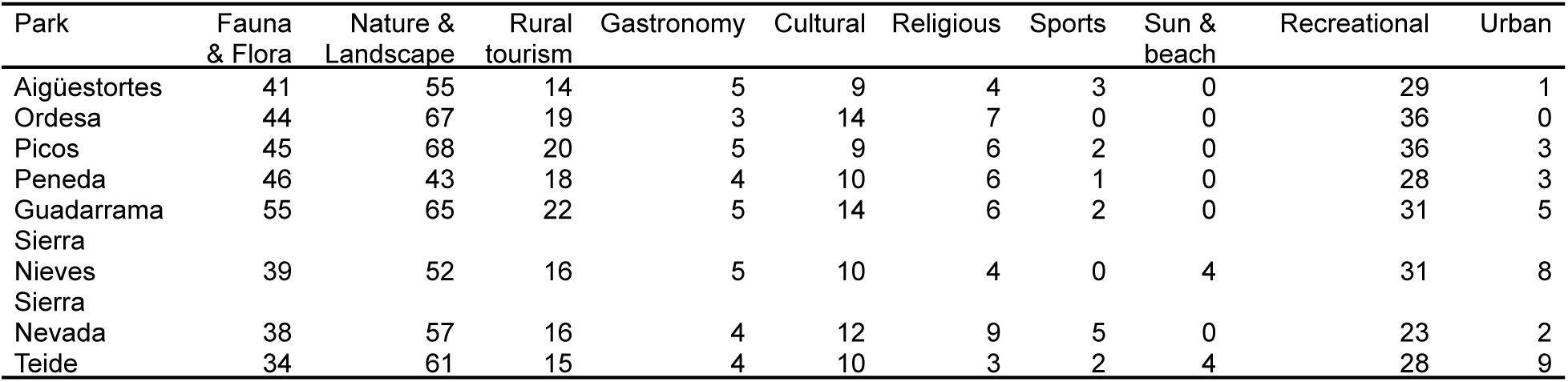
Number of labeled images within the 10 CES-related classes in each park included in the independent evaluation dataset. The total number of images differs from 1082, as each image could be assigned up to three CES categories from the 10 CES-related classes. Images classified as non-relevant were excluded.

| Park | Fauna & Flora | Nature & Landscape | Rural tourism | Gastronomy | Cultural | Religious | Sports | Sun & beach | Recreational | Urban |
| --- | --- | --- | --- | --- | --- | --- | --- | --- | --- | --- |
| Aigüestortes | 41 | 55 | 14 | 5 | 9 | 4 | 3 | 0 | 29 | 1 |
| Ordesa | 44 | 67 | 19 | 3 | 14 | 7 | 0 | 0 | 36 | 0 |
| Picos | 45 | 68 | 20 | 5 | 9 | 6 | 2 | 0 | 36 | 3 |
| Peneda | 46 | 43 | 18 | 4 | 10 | 6 | 1 | 0 | 28 | 3 |
| Guadarrama | 55 | 65 | 22 | 5 | 14 | 6 | 2 | 0 | 31 | 5 |
| Sierra Nieves | 39 | 52 | 16 | 5 | 10 | 4 | 0 | 4 | 31 | 8 |
| Sierra Nevada | 38 | 57 | 16 | 4 | 12 | 9 | 5 | 0 | 23 | 2 |
| Teide | 34 | 61 | 15 | 4 | 10 | 3 | 2 | 4 | 28 | 9 |

To mitigate spatial and content bias, we implemented a selection process similar, but more restrictive, to the well-known “Photo-User-Days” (PUD) approach. Specifically, within each cell of the 100 m, we retained only one photo per user and label. This approach ensures that label consensus within each cell is driven by the agreement among different users, thereby reducing individual user bias. Thus, after excluding those images selected for independent evaluation and applying the described filter, a total of 67,196 labeled images remained for the CES modeling process (see Table 2).

**Table 2.**
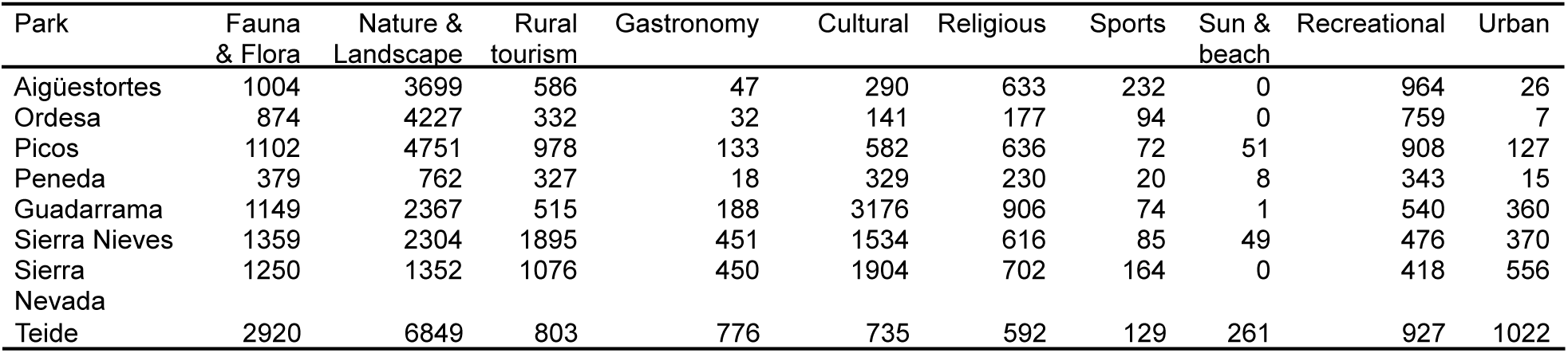
Number of labeled images within the 10 CES-related classes in each park and selected for CES modeling (presences).

### 2.3. Spatial predictors

To identify the spatial predictors influencing the provision and distribution of CES, a systematic literature review of studies on spatial modeling of CES published up to December 31, 2024, was conducted using the Web of Science (WOS) and Scopus databases, which identified a total of 362 predictors used in CES modeling (Navarro et al, 2025). Based on this review, we selected a set of 297 predictors categorized into 7 groups. These groups encompass accessibility, climate, ecosystem structure, ecosystem functioning, geodiversity, land use and land cover, and tourism and culture related variables. Given the limited expert knowledge on the specific factors influencing the provision of CES, this approach ensures a comprehensive representation of the potential drivers of CES provision. All predictors were cropped to fit the study area, reprojected to ETRS89-extended / LAEA Europe (EPSG: 3035), and resampled to a 100 m resolution.

To reduce the dimensionality, we employed a two-step “embedded” covariate selection procedure using the covsel R package (Adde et al., 2023). In the first step, variables with a Pearson correlation coefficient > |0.7| were excluded through a sequential process conducted at three levels: the variable level, the group level, and across all remaining predictors. In the second step, the filtered predictors were used to train generalized linear models (GLM) and random forest (RF) models, which ranked predictors based on their importance, using the maximum absolute values of the regularized regression coefficients for GLM and the Mean Decrease Gini index for RF. The final ranking was determined by summing the ranks from both models. Finally, the number of selected predictors was dependent on the number of presences of the target CES, ranging from the highest integer of the logarithm (base 2) of the number of occurrences up to a maximum of 12, thus limiting the complexity of the models (Brun et al., 2020; Sillero et al., 2021).

### 2.4. CES modelling

The maximum entropy (MaxEnt) approach (Phillips et al., 2006) was employed to model CES using the biomod2 (version 4.2-4) R package (Thuiller et al., 2023). MaxEnt is a non-stochastic presence-only method (i.e., results do not vary each time it is run), which is well-suited for handling low presences sample sizes, deal with both continuous and categorical predictors, and has demonstrated strong predictive performance while maintaining computational efficiency (Sillero et al., 2021; Valavi et al., 2022). Since MaxEnt is sensitive to sampling bias, we generated 10 pseudo-absence replicates following the “target-group background” approach (Phillips et al., 2009). This method estimates sampling effort for the target CES by leveraging the occurrences of similar CES, assuming that the sampling effort is the same across all. The sampling intensity is then converted into a continuous probability surface, which is used to weight the selection of background points, thereby restricting the available background to a specified distance from records of similar CES. As presences vary by National Park, we generated the target-group background using all CES presences at the National Park level. Finally, to deal with class-imbalance problems, each target-group replicate selects the same number of pseudo-absences as presences (Liu et al., 2009).

The predictive performance and discrimination ability of MaxEnt models was assessed with the validation data set through (i) the Area Under the Receiver Operating Characteristics (AUC-ROC) curve and (ii) True-skill statistic (TSS). AUC-ROC quantifies the model’s ability to distinguish between presences and absences across different threshold values. Values range from 0 (where the prediction is 100% incorrect) to 1 (perfect discrimination), with higher values indicating better predictive performance (Lobo et al., 2008). TSS, evaluates model accuracy by incorporating both sensitivity (true positive rate) and specificity (true negative rate). It ranges from -1 to 1, where values close to 1 indicate high predictive power, and values close to or less than zero indicating performance no better than random expectation (Allouche et al., 2006). To minimise uncertainties evaluating the models, a 10-fold cross-validation procedure was implemented, partitioning the dataset into ten equally sized folds. In each iteration, nine folds were used for model training, while the remaining fold served for validation. Given the 10 pseudo-absence replicates and the 10-fold cross-validation, each model was run 100 times. Since AUC-ROC is a robust threshold-independent measure of a model’s ability to discriminate presence from absence, the ensemble models were then built using any model runs with AUC-ROC scores of > 0.6. These models were assembled using the median, while uncertainty was quantified by calculating the coefficient of variation across the models.

Following this methodology, we developed models using two different approaches. In the first approach, models were fitted for each CES label irrespective of the study area (i.e, all occurrences of a given CES label were used regardless of the park) and then separately projected for each study area. This approach assumes that people’s choices and preferences for a given CES are consistent across different locations. In the second approach, models were developed and projected separately for each CES label within each study area, assuming that people’s choices and preferences for a given CES are influenced by the park’s specific environmental, landscape, and biophysical context.

### 2.5. Independent CES models evaluation

Models of both approaches were evaluated at study area level using the Boyce index as in Hirzel et al., (2006) with the independent and manually labeled subset of 1082 images. The Boyce index is the only evaluation metric specifically designed for presence-only based predictions and measures how much model predictions differ from a random distribution of observed presences (Boyce et al., 2002). Its values range between -1 and 1, where positive values indicate that predicted presences align well with the distribution of presences in the evaluation dataset, suggesting good model performance. Values close to zero imply that the model’s predictions are not statistically different from a random distribution, while negative values indicate counter predictions, meaning the model predicts lower suitability in areas where presences are more frequent (Hirzel et al., 2006; Sillero et al., 2021). Therefore, in addition to evaluating the model itself, for each CES, the model with the highest Boyce value between the two approaches will indicate whether people’s preferences and choices are primarily influenced by the park’s specific environmental, landscape, and biophysical context or whether they remain consistent regardless of location.

Boyce index was calculated from the model predictions using the ecospat R package (Broennimann et al., 2025). Additionally, a mean comparison between Boyce values of both approaches were performed with Wilcoxon-Mann–Whitney test (at α = 0.1) to reveal statistically significant differences at CES level. Thus, the CES models from the approach that best explains people’s choices and preferences will be selected to identify the most informative predictors for each CES, spatially project the CES in each park, and guide further analysis. Of that, individual models with a Boyce index below 0.5 were also excluded.

### 2.6. Statistical analysis over CES projections

To evaluate the influence of protection levels on both CES suitability and richness, we used CES models derived from the best-performing modeling approach for each CES category. For each combination of protection levels, we extracted 5,000 pixel values via bootstrapping, repeating the procedure 100 times to ensure robustness. Subsequently, we developed two models: a generalized linear model (GLM) for CES suitability and a linear model (LM) for CES richness.

First, we fitted a GLM for each CES category, using predicted suitability as the response variable and protection level and study area as explanatory variables. Given that CES suitability values are bounded between 0 and 1, a beta distribution was applied. This model enabled us to examine whether protected areas generally support higher suitability for certain CES categories and whether these effects differ among CES types.

Second, we applied a linear model to assess CES richness, calculated as the sum of ROC-filtered continuous suitability values across all CES categories. To allow for comparability among study areas, CES richness was normalized by dividing by the number of reliable CES categories, yielding a standardized value between 0 and 1. This analysis aimed to determine whether stricter protection regimes are associated with increased or decreased CES richness.

## 3. Results

### 3.1. Performance of deep learning-based Social Media data to map CES

Most CES categories exhibited positive Boyce values higher than 0.5, indicating a good model performance (Figure 2; see Figures S1 y S2 for ROC and TSS validation assessment with 10-folds cross validation). The All Parks approach yielded relatively consistent Boyce values across all parks, whereas the By Park approach showed greater variations, reaching for some parks in particular values far away from their median values.

**Figure 2.**
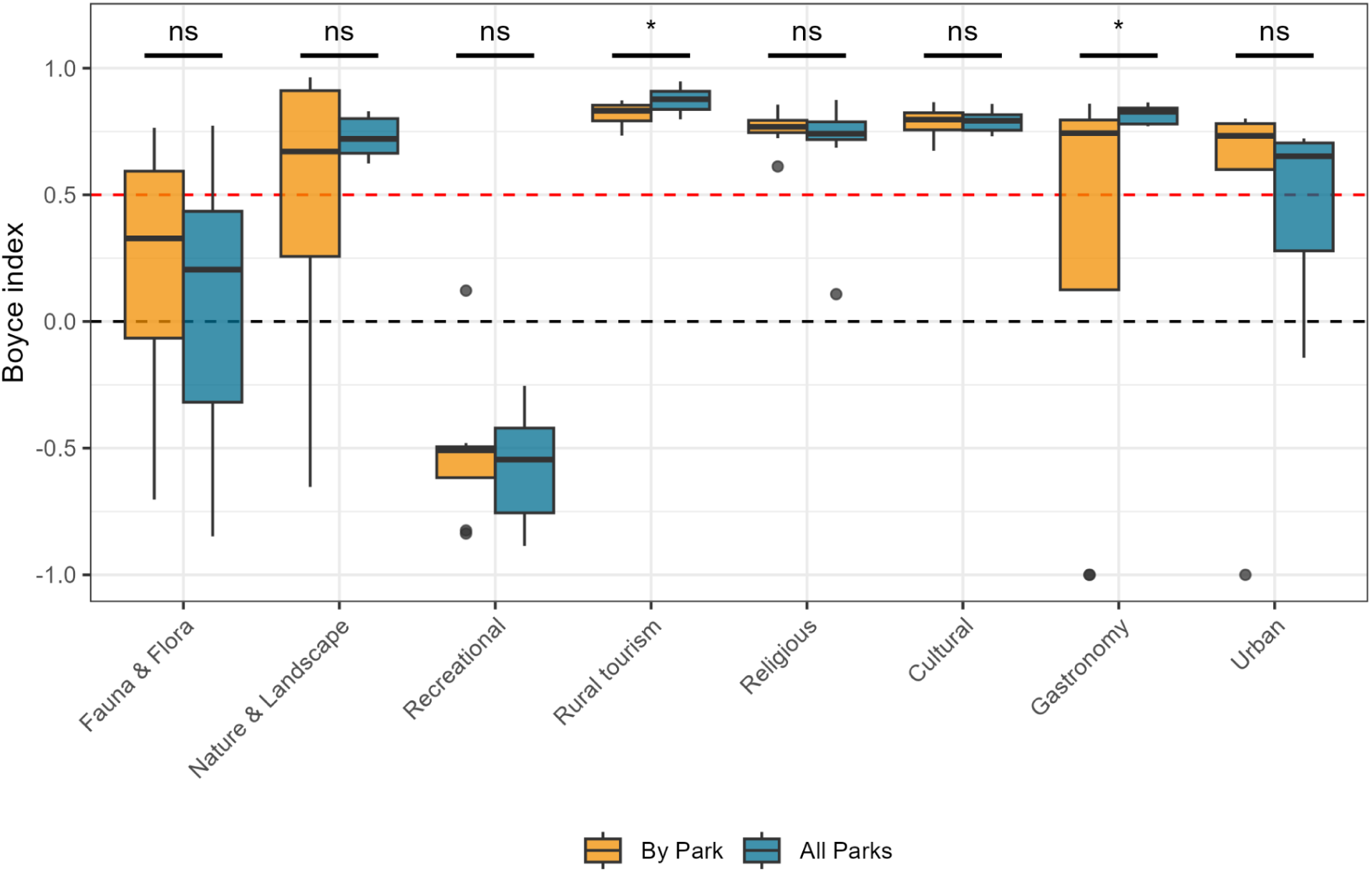
Boyce index distribution for the different CES and approaches shown as box-and-whisker plots (midline: median; boxes: 25% and 75% quartiles; whiskers: largest values no further than 1.5 times the interquartile range; and dots: outlier values), and statistical significance of differences between approaches indicated above each category using Wilcoxon-Mann–Whitney test: ns = not significant, * = p < 0.10, ** = p < 0.05, *** = p < 0.01.

When comparing both approaches, the All Parks modeling approach resulted in higher performance for Nature & Landscape, Rural tourism and Gastronomy, while the By Park approach outperformed in Religious, Cultural, Urban and Flora & Fauna, although in the latter only for Sierra Nevada and Teide. It is noteworthy that in most cases, the median values for both approaches when they are able to predict well are very close (see for example the Rural tourism, Religious, or Cultural categories). Recreational showed median values close to -1 in both approaches, indicating that model predictions did not align with the observed presence distribution, while the Fauna & Flora CES had, in general, Boyce values below 0.5, suggesting low model performance.

For each CES, the model derived from the most successful methodological approach was projected onto the geographical space of each study area, generating the suitability maps (Figure 3). Overall, the highest suitability areas (represented in red) for Fauna & Flora were located outside the National Park boundaries. The distribution of Nature & Landscape CES was more strongly associated with the boundaries of national parks, particularly in Sierra Nevada and Teide. While the highest suitability values were generally concentrated within protected areas, considerable suitability was also observed beyond park limits, especially in coastal zones. Suitability for Rural Tourism and Gastronomy CES showed a strong spatial association with human settlements. In coastal study areas, Gastronomy CES also demonstrated a pronounced affinity for littoral zones. Cultural, and Religious CES were also predominantly associated with human settlements and culturally significant landmarks. A particular example is the Santuario de Nuestra Señora de Covadonga in Picos de Europa, which emerged as a focal point for religious and cultural CES. Similarly, Rural tourism, Gastronomy and Urban CES displayed a spatial association with human settlements.

**Figure 3.**
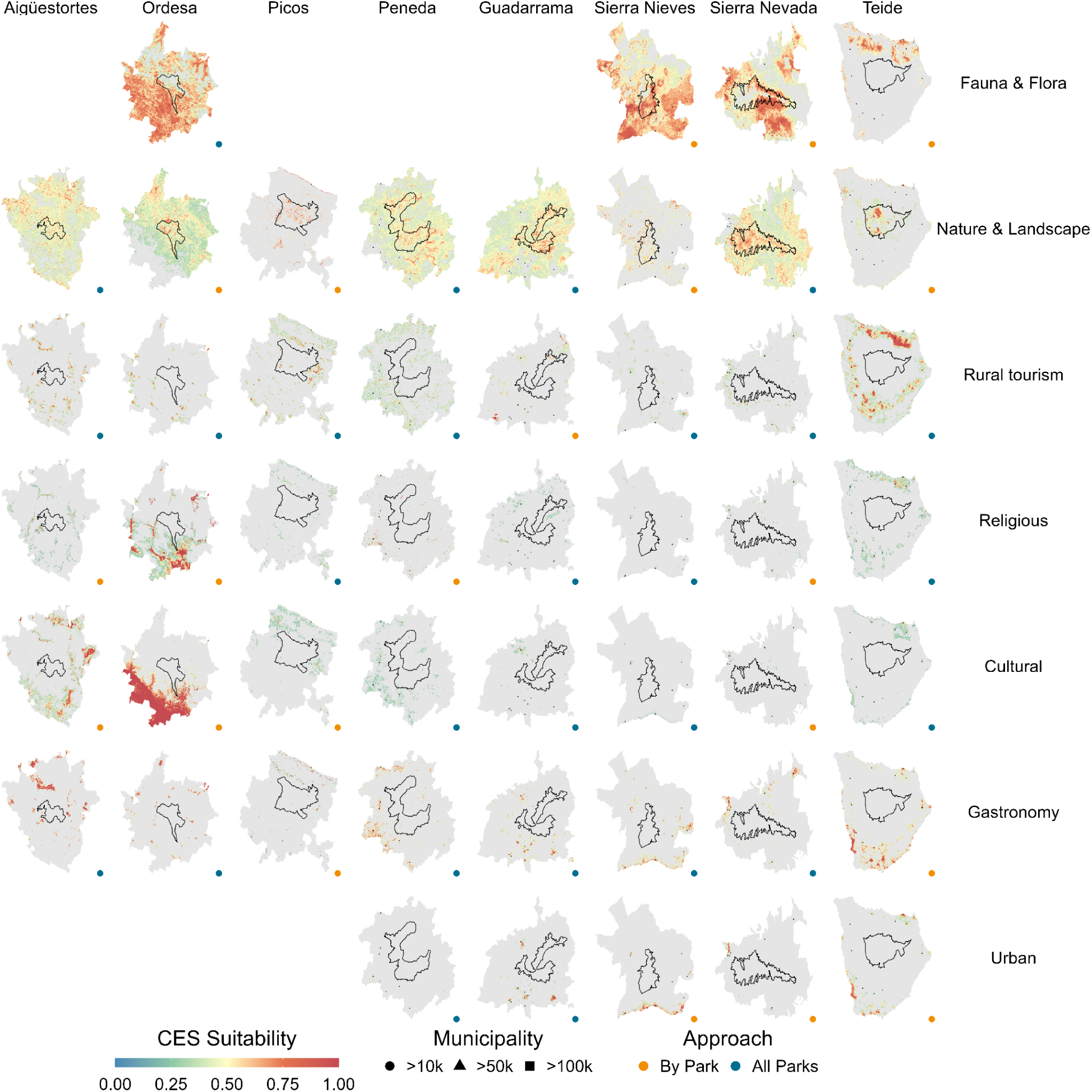
CES suitability provided within each study area using people’s choices and preferences as a proxy. The black line delineates the National Park boundaries. Suitability values below the ROC threshold were filtered. The municipalities represent the population centres with more than 10,000 inhabitants classified by their population census, obtained from the National Statistical Institutes of Spain and Portugal.

### 3.2. Determinants of CES preferences

Overall, the most representative variable groups selected for By Park approach were Tourism and Culture, Ecosystem functioning and Climate, while Climate, Ecosystem functioning and Ecosystem structure were the most selected for All parks approach. However, only a limited number of variables contributed significantly to the models. Focussing only on the three most important variables for each model (Figures 4 and 5), the comparison of variable importance between the two modeling approaches revealed that, while some key predictors are shared between the ‘All Parks’ model and the individual park models, there are notable differences in both the identity and ranking of the most influential variables.

**Figure 4.**
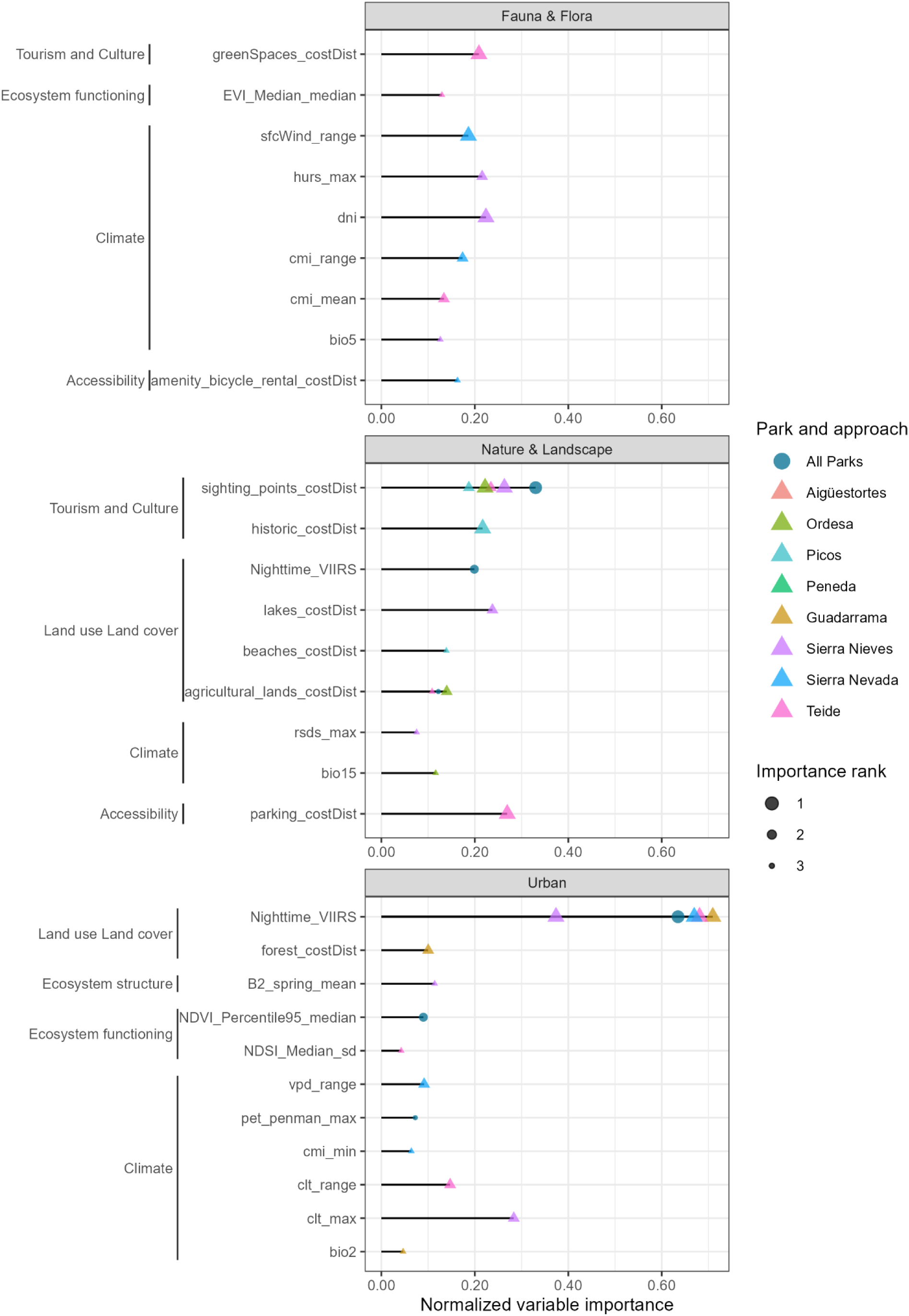
Variable importance for the Fauna & Flora, Nature & Landscape, Urban CES categories, based on two modeling approaches: a global model that combines data from all parks (circles, “All Parks”) and park-specific models developed individually for each site (triangles, one per park). The importance rank reflects whether a variable was the first, second, or third most influential in model performance. To the left of each variable, its associated thematic group (e.g., Ecosystem functioning, Ecosystem structure, etc…) is displayed to aid in the interpretation of predictor relevance. CES categories with a Boyce index below 0.5 were excluded due to limited predictive reliability.

**Figure 5.**
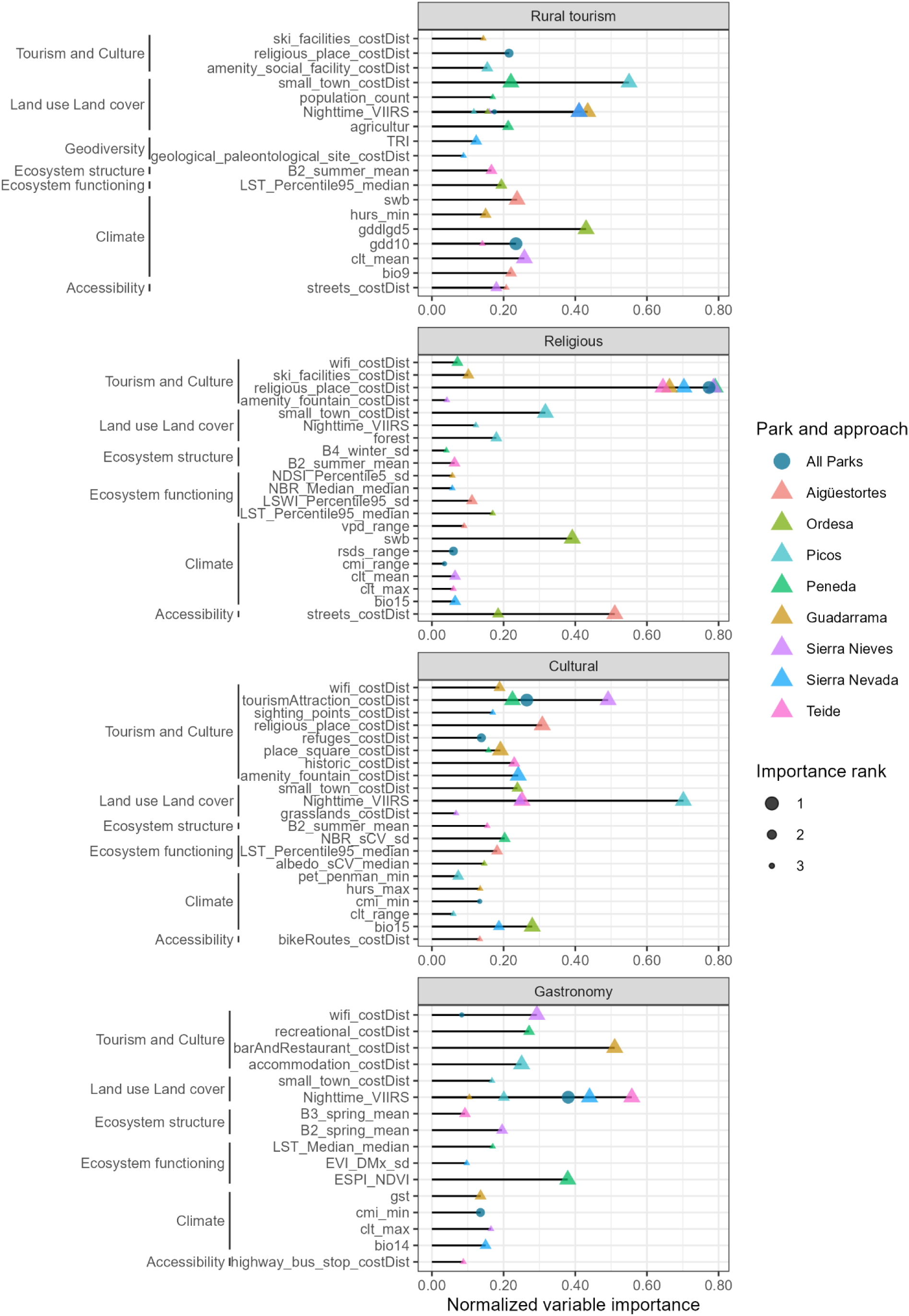
Variable importance for the Rural tourism, Gastronomy, Cultural and Religious CES categories, based on two modeling approaches: a global model that combines data from all parks (circles, “All Parks”) and park-specific models developed individually for each site (triangles, one per park). The importance rank reflects whether a variable was the first, second, or third most influential in model performance. To the left of each variable, its associated thematic group (e.g., Ecosystem functioning, Ecosystem structure, etc…) is displayed to aid in the interpretation of predictor relevance. CES categories with a Boyce index below 0.5 were excluded due to limited predictive reliability.

For Fauna & Flora, none of the variables was selected by the three park-specific models. For Nature & Landscape, there was strong agreement across all models, with distance to sighting points consistently identified as the most important predictor, though secondary variables differed between parks. A similar pattern was observed for Urban and Gastronomy, where the presence of nighttime lights was consistently identified as the primary predictor across all models. In the case of Gastronomy, distance to bars and restaurants, accommodation sites and wifi appeared as influential variables for park-specific models.

For Rural Tourism, nighttime lights emerged as the most influential predictor in two of the eight study areas and ranked as the third most important variable in several other models. Distance to small towns was identified as the top predictor in two park-specific models, while the global model (All parks) and the remaining park-specific models highlighted different variables as most important. A similar pattern was observed for Cultural, where variables such as proximity to tourist attractions and nighttime lights frequently appeared among the most influential across parks and modeling approaches, though no clear consensus emerged regarding a dominant set of predictors. Lastly, for Religious CES, distance to religious sites was the most important predictor in both the global model and in five park-specific models, with normalized importance scores substantially higher than those of any other variable, though there was little agreement on which variables were of lesser relevance.

### 3.3. Influence of protection level and parks over CES suitability

All levels of protection had a positive effect CES suitability (Figure 6). In general, areas under some form of protection exhibited higher suitability values than unprotected areas (000). Among these, Biosphere Reserves (001) consistently showed relatively high and stable suitability across all CES types. Protection levels that included National Parks and Natura 2000 sites (e.g., 010, 110, 111) were particularly effective in enhancing suitability for Fauna & Flora and Nature & Landscape CES. In contrast, CES associated with Urban, Gastronomy, and Rural Tourism exhibited lower suitability values across all protection levels, including those with national park and Natura 2000 designations. Although the suitability of these services increased slightly under protection, the improvements were modest and remained below those observed for nature-based services.

**Figure 6.**
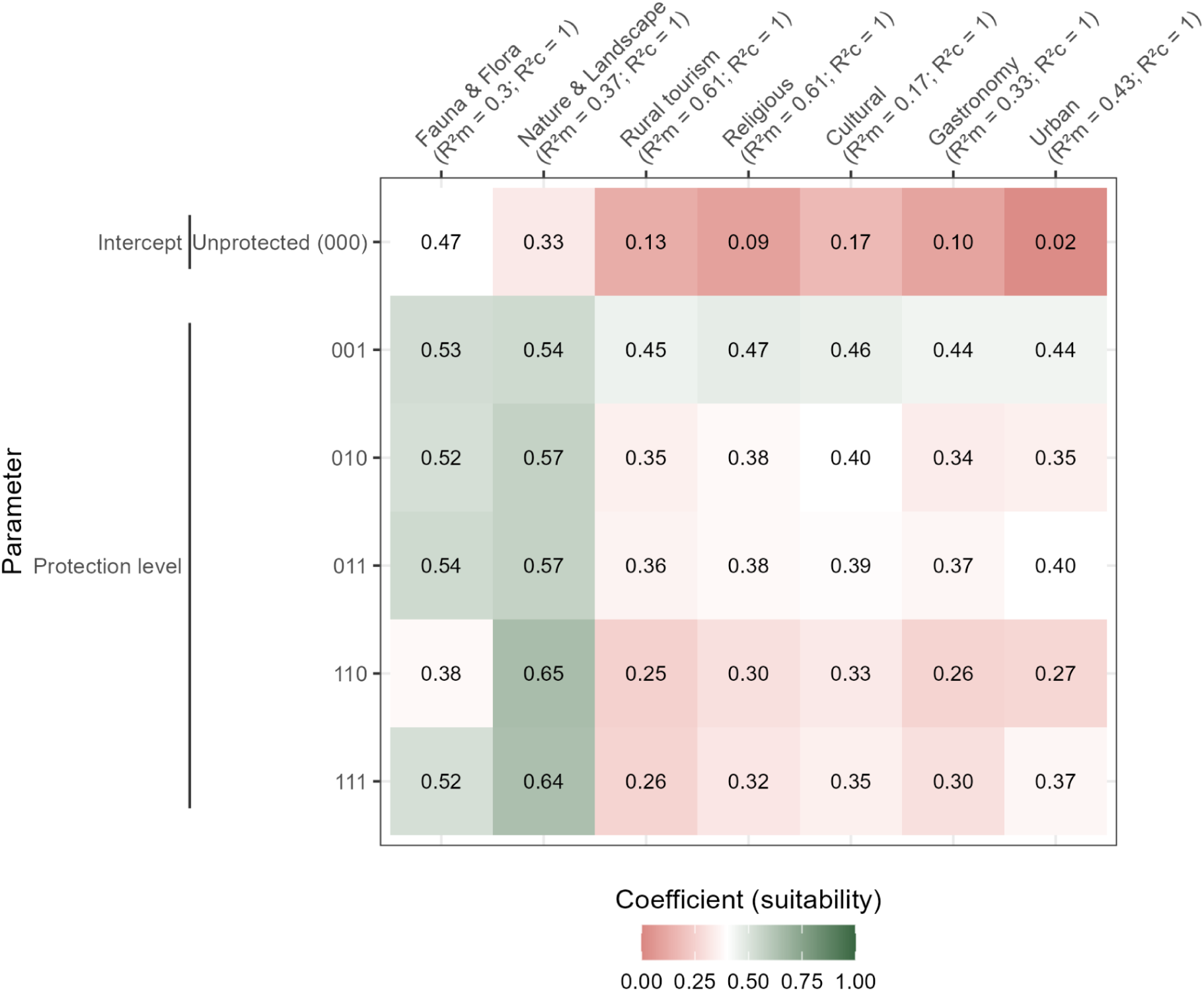
Summary of GLMs coefficients evaluating the effects of protection status. The models assume a beta distribution for the response variable. Coefficients were transformed from log-odds scale to suitability using the inverse-logit transformation: Suitability = 1 / (1 + exp(−coefficient)). Only coefficients that are statistically significant (p ≤ 0.05) are highlighted in color.

## 4. Discussion

Ongoing…

## 5. Conclusion

Ongoing…

## Acknowledgement

We thank Manuel Merino-Ceballos and Andrea Ros-Candeira for their support in the download, preparation and labelling of the data. JML received credits from the OpenAI Researcher Access Program.

