## Supplementary material for "What drives cultural ecosystem services in mountain protected areas? An AI-assisted answer using social media"

Table S1. Percentage (%) of each protection level by park. The first position of the protection code indicates whether the cell is within a national park, the second denotes inclusion within a Natura 2000 site, and the third specifies whether the cell falls within a biosphere reserve. Presence and absence of protection were represented by ones (1) and zeros (0), respectively. For example, '110' indicates that the cell is protected under both a national park and a Natura 2000 site but is not part of a biosphere reserve.

| Protection code | Aigüestortes | Ordesa | Picos | Peneda | Guadarrama | Sierra Nieves | Sierra Nevada | Teide |
|---|---|---|---|---|---|---|---|---|
| 000 | 42.7 | 23.1 | 35.8 | 43.2 | 29.5 | 48.4 | 51.8 | 49.0 |
| 001 | | 14.1 | 2.0 | 27.9 | 7.4 | 22.8 | | |
| 010 | 52.8 | 37.2 | 37.4 | 2.8 | 29.4 | 1.5 | 18.2 | 39.0 |
| 011 | | 19.2 | 11.4 | 11.5 | 22.9 | 20.8 | 14.8 | |
| 110 | 4.5 | | 0.5 | | 1.9 | | | 12.1 |
| 111 | | 6.3 | 12.9 | 14.7 | 8.8 | 7.3 | 15.2 | |

Table S2. Description of the type of images included in each CES-related category.

| CES label | Description |
|---|---|
| Nature & Landscape | Images whose main theme is nature in general, including landscapes. Captured in a wide-angle shot. |
| Fauna & Flora | Images whose main theme is fauna and/or flora or their elements. Captured in close-up or mid-range shots. |
| Recreational | Images whose main theme is the use of recreational areas (scenes of people spending time in recreational spaces or similar public infrastructures). Captured in close-up, mid-range, or wide-angle shots. |
| Sports | Images whose main theme is sports activities and/or their elements. Captured in close-up or wide-angle shots. |
| Cultural | Images whose main theme is cultural elements (e.g., livestock movement, a traditional loom, activities such as basket weaving, photographs of Alpujarra-style woven textiles, etc.). Captured in close-up or wide-angle shots. |
| Religious | Images whose main theme is religious elements (e.g., Virgin Mary, processions, pilgrimages, churches). Captured in close-up or wide-angle shots. |
| Gastronomy | Images whose main theme is gastronomy (e.g., dinner in a restaurant, traditional food products). Captured in close-up or wide-angle shots. |
| Rural tourism | Images whose main theme is rural tourism elements (e.g., rural accommodations, rural villages, small towns, or hamlets). Captured in close-up, mid-range, or wide-angle shots. |
| Urban | Images whose main theme is urban elements (e.g., houses, streets, parks). Captured in close-up or wide-angle shots. |
| Sun & beach | Images whose main theme is sun-and-beach tourism elements (e.g., photographs of people or events taking place on the beach in a leisure context). |

Table S3. F1-score for each CES category based on the manually and independently annotated random subset of 1082 images covering all parks, municipalities, and labels.

| CES | F1-score |
|---|---|
| Cultural | 0.76 |
| Fauna & Flora | 0.94 |
| Gastronomy | 0.94 |
| Nature & Landscape | 0.92 |
| Not relevant | 0.81 |
| Recreational | 0.85 |
| Religious | 0.99 |
| Rural tourism | 0.83 |
| Sports | 0.25 |
| Sun & beach | - |
| Urban | 0.77 |

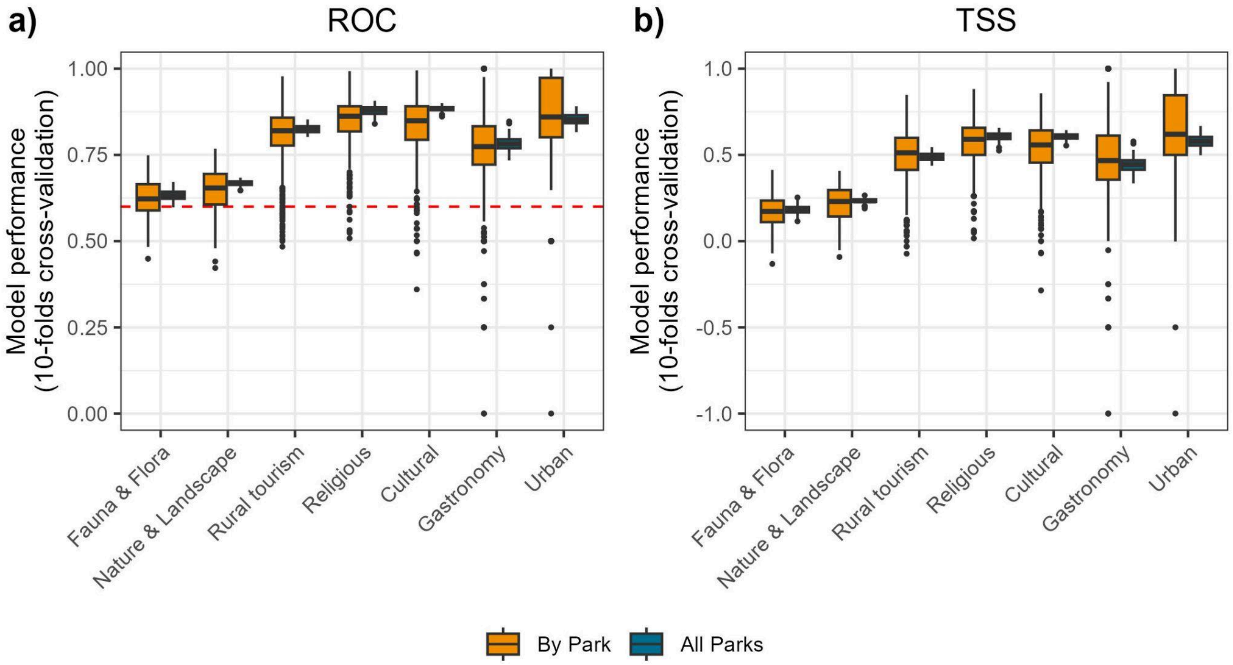


Figure S1. Model performance by CES and approach using the a) ROC and b) TSS as evaluation metric. The dashed red line indicates the ROC threshold considered to build ensemble models.

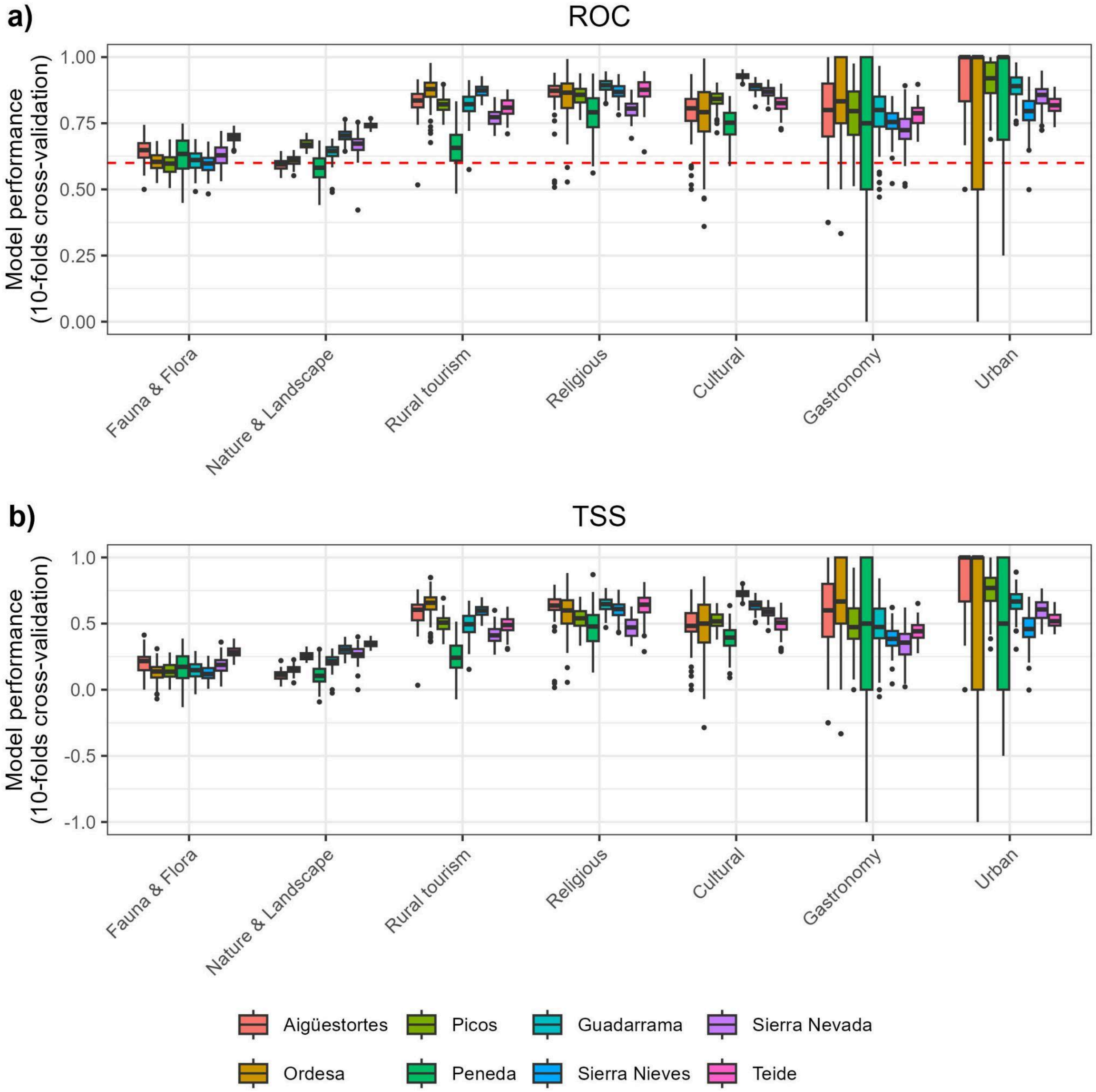


Figure S2. Model performance by CES and park for By Park approach using the a) ROC and b) TSS as evaluation metric. The dashed red line indicates the ROC threshold considered to build ensemble models.
